# Multi-monoubiquitination controls VASP-mediated actin dynamics

**DOI:** 10.1101/2023.07.16.549237

**Authors:** Laura E. McCormick, Cristian Suarez, Laura E. Herring, Kevin S. Cannon, David R. Kovar, Nicholas G. Brown, Stephanie L. Gupton

## Abstract

The actin cytoskeleton performs multiple cellular functions, and as such, actin polymerization must be tightly regulated. We previously demonstrated that reversible, non-degradative ubiquitination regulates the function of the actin polymerase VASP in developing neurons. However, the underlying mechanism of how ubiquitination impacts VASP activity was unknown. Here we show that mimicking multi-monoubiquitination of VASP at K240 and K286 negatively regulates VASP interactions with actin. Using in vitro biochemical assays, we demonstrate the reduced ability of multi-monoubiquitinated VASP to bind, bundle, and elongate actin filaments. However, multi-monoubiquitinated VASP maintained the ability to bind and protect barbed ends from capping protein. Lastly, we demonstrate the introduction of recombinant multi-monoubiquitinated VASP protein altered cell spreading morphology. Collectively, these results suggest a mechanism in which ubiquitination controls VASP-mediated actin dynamics.

## Introduction

The dynamic remodeling of the actin cytoskeleton regulates many aspects of cell biology, including cell division, morphology, motility, and endocytic trafficking. Starting from single monomers, actin polymerizes into filaments that interact with a myriad actin regulatory proteins to form a variety of different cytoskeletal architectures (Pollard and Cooper, 2009). Local actin reorganization, polymerization, and depolymerization are dependent upon spatiotemporal regulation by these different proteins.

Amidst the abundance of actin regulatory proteins, the Ena/VASP protein family (Mena, VASP, and EVL) occupies a specialized niche (Krause et al., 2003; Bear and Gertler, 2009). The tetrameric Ena/VASP proteins bind and bundle parallel actin filaments. They also bind to actin monomers and the fast growing “barbed” ends of actin filaments, where they enhance the addition of monomers and protect the barbed end from capping protein, which terminates actin polymerization (Faix and Rottner, 2022; Krause et al., 2003). The combination of these functions position Ena/VASP proteins as multi-faceted regulators of filopodia—finger-like bundled actin-rich protrusions that extend from the cell periphery. Although their precise function varies by cell type, filopodia are considered sensors that explore the local environment, influencing cell migration and morphology changes (Gupton and Gertler, 2010; Jacquemet et al., 2015). For example, developing murine cortical neurons require Ena/VASP proteins for filopodia formation and subsequent neurite initiation (Dent et al., 2007; Kwiatkowski et al., 2007). Impairment of Ena/VASP activity after neurite initiation disrupts growth cone filopodia formation and response to guidance cues (Lebrand et al., 2004). Lastly, VASP and EVL play a role in dendritic filopodia and dendritic spine formation (Lin et al., 2010; Parker et al., 2023).

Previously, we identified the brain-enriched E3 ubiquitin ligase TRIM9 as an interaction partner of VASP (Menon et al., 2015, 2021). VASP and TRIM9 colocalized at the tips of growth cone filopodia in developing neurons, suggesting TRIM9 also regulates these structures. Loss of Trim9 increased filopodia number and stability in the growth cone and impaired axon pathfinding. We also showed that TRIM9 was required for the non-degradative ubiquitination of VASP in neurons (Menon et al., 2015). Interestingly, this was a reversible modification; addition of the guidance cue netrin-1 resulted in VASP deubiquitination and a consequential increase in filopodia stability and number. Furthermore, experiments utilizing ubiquitin mutant constructs incapable of ubiquitin chain formation suggested VASP was mono-or multi-monoubiquitinated, as opposed to polyubiquitinated (Boyer et al., 2020). Although K48-linked ubiquitination is associated with proteasomal degradation, monoubiquitination is appreciated to regulate protein localization and activity (Baker et al., 2013; Lin et al., 2016; Nakagawa and Nakayama, 2015). As VASP ubiquitination reduced filopodial stability, we hypothesized that mono or multi-monoubiquitination negatively regulated VASP activity.

In this study, we investigate the mechanistic impact of ubiquitination on VASP function. We identified two prominent ubiquitination sites within VASP. Through biochemical reconstitutions, we demonstrate that mimicking multi-monoubiquitination of VASP reduces actin filament binding and bundling. Furthermore, both mono- and multi-monoubiquitination diminishes the ability of VASP to enhance actin filament elongation. Our results suggest that ubiquitinated VASP maintains barbed end association but is unable to efficiently add new actin monomers to the filament. In murine embryonic fibroblasts, electroporation of multi-monoubiquitinated VASP alters cell spreading morphology and decreases localization to the lamellipodia and filopodial tip. Collectively, our findings mechanistically describe the negative regulation of VASP activity by reversible mono- and multi-monoubiquitination.

## Results

### Identification of ubiquitinated lysine residues on VASP

We previously demonstrated that non-degradative ubiquitination of VASP impacted filopodial dynamics and axon guidance (Menon et al., 2015). Subsequent studies indicated that VASP was likely mono-or multi-monoubiquitinated (Boyer et al., 2020). However, the mechanistic consequence of VASP ubiquitination—and how this impacted filopodial dynamics—is unknown. Consistent with neurons, VASP ubiquitination in HEK293 cells was dependent upon the E3 ubiquitin ligase TRIM9 (Menon et al., 2015). To determine which residues on VASP were ubiquitinated, we immunoprecipitated exogenous Myc-VASP from HEK293 cells under denaturing conditions. Following separation by SDS-PAGE and immunoblotting, we observed two identifiable VASP bands: a prominent, unmodified VASP band at approximately 65 kD, and a higher molecular weight VASP band (75 kD) that comigrated with ubiquitin, indicating this was ubiquitinated VASP (VASP-Ub, Fig 1A). This shift in molecular weight and relative ratio of unmodified and ubiquitinated VASP is similar to what we have previously observed with Myc-VASP in HEK293 cells and endogenous VASP in murine cortical neurons (Menon et al., 2015; Boyer et al., 2020).

**Fig. 1.**
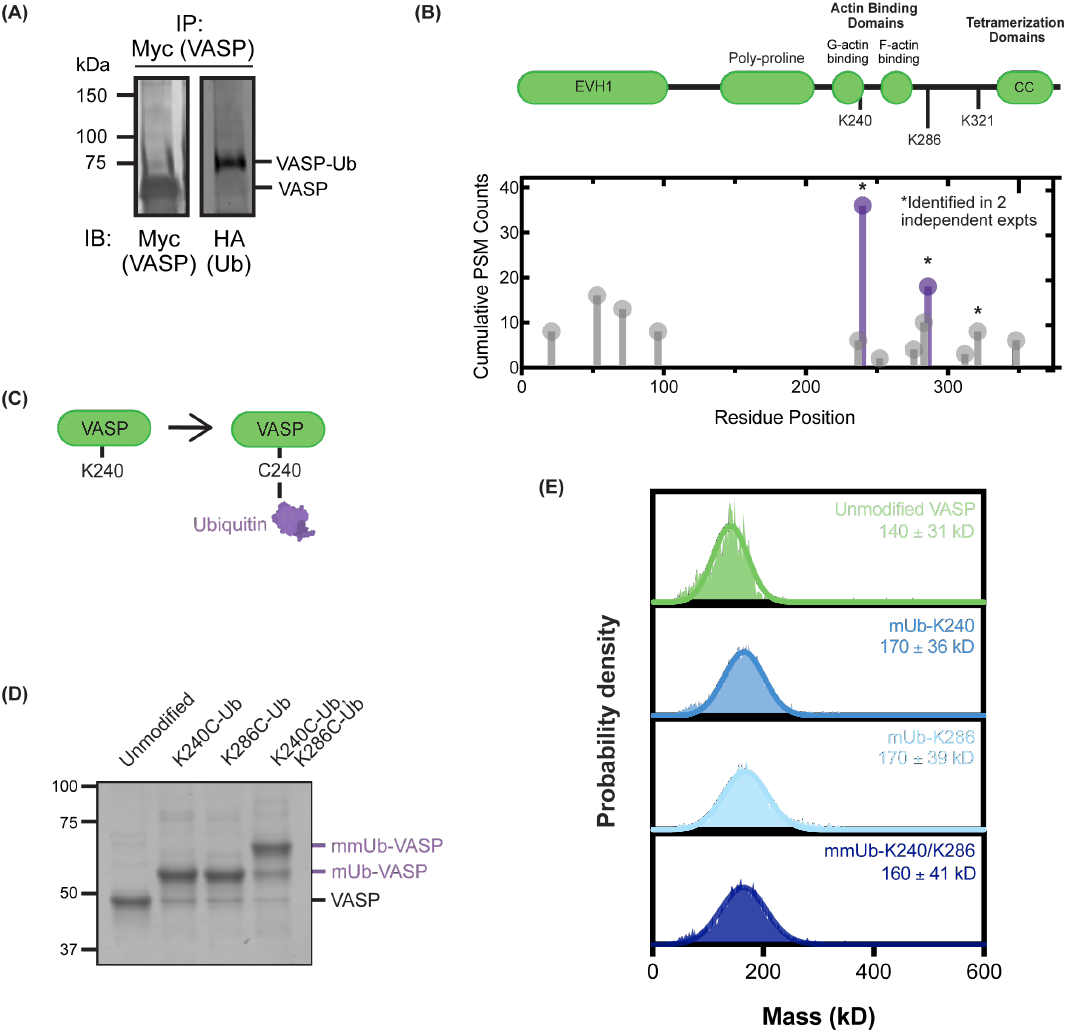
VASP is ubiquitinated at K240 and K286. (A) Western blot of ubiquitinated Myc-VASP. HEK293 cells were transfected with Myc-VASP and HA-Ubiquitin, lysed, and boiled in denaturing buffer before Myc immunoprecipitation. (B) VASP domain architecture and proteomic detection of ubiquitination sites on VASP by mass spectrometry. Purple residues were selected for follow-up study due to their abundance and replicability. (C) Chemical ubiquitination scheme, demonstrating the bismaleimidoethane-mediated crosslinking of ubiquitin G75C to Cysteine-240 on VASP. (D) Coomassie gel of purified, ubiquitinated VASP at indicated locations. (E) Mass photometry distributions of unmodified and ubiquitinated VASP. The estimated molecular weight ± SD (rounded to two significant figures) was calculated with a Gaussian curve fit. These graphs show one representative trace for each construct (N=2).

Immunoprecipitated VASP was separated on a SDS-PAGE gel and excised for analysis by mass spectrometry. Following enzymatic digestion and analysis, we identified numerous lysine residues as ubiquitination sites on VASP by unbiased mass spectrometry and confirmed by targeted analysis (Fig 1B). Of multiple lysine residues identified, three ubiquitination sites replicated across independent experiments and different enzymatic digestions. From these three sites, we selected the two most abundant ubiquitination sites, K240 and K286, for investigation (Fig S1). K240 is located within the monomeric actin binding domain (GAB) of VASP and K286 is located near to the filamentous actin binding domain (FAB).

### Generation of purified, ubiquitinated VASP mimics

To investigate how ubiquitination at these sites altered protein function, we purified and ubiquitinated recombinant human VASP in vitro. Unlike phosphorylation or acetylation, modification of a protein with ubiquitin, an 8 kDa protein, cannot be mimicked with a single amino acid substitution. Instead, we utilized a chemical ubiquitination approach, in which a modified recombinant ubiquitin (Ubiquitin1-74, G75C) was covalently conjugated to available cysteine residues on VASP (Fig 1C). Three endogenous cysteines are present in human VASP, requiring their modification to specifically ubiquitinate residues of interest. Fortuitously, a cysteine-light version of VASP (VASPCCC-SSA) was previously generated and shown to have identical actin binding and elongation activity to unmodified VASP (Hansen and Mullins, 2010).

Previously, we showed VASP was monoubiquitinated or multi-monoubiquitinated (Boyer et al., 2020). As phosphorylation of VASP at Ser157 causes a disproportionally large apparent molecular weight shift (4kDa) (Smolenski et al., 1998), we could not interpret the apparent molecular weight shift observed for VASP-Ub by SDS-PAGE as mono or multi-monoubiquitination. Therefore, we created two constructs with single cysteine mutations—VASP K240C and VASP K286C—within the cysteine-light VASP background construct to mimic two versions of monoubiquitinated VASP (mUb-K240 and mUb-K286). We also created a double cysteine mutant VASP K240C, K286C to mimic a multi-monoubiquitinated version of VASP protein (mmUb-K240/K286) (Fig 1D). With each of these recombinant proteins, we achieved over 90% modification with ubiquitin after chemical crosslinking. Following successful chemical ubiquitination, we used the bacterial sortase enzyme to conjugate a fluorophore labeled peptide, TAMRA-PEG6-LPETGG, to a poly-glycine motif added at the N-terminus of VASP.

### VASP-Ub does not disrupt tetramerization

The COOH-terminal end of VASP contains an essential tetramerization domain (Fig 1B). Studies manipulating oligomerization demonstrated Ena/VASP oligomer number directly correlated with the actin elongation rate in vitro (Brühmann et al., 2017) and filopodia formation in cultured cells (Harker et al., 2019). As VASP-Ub was associated with impaired filopodia number and stability in cultured neurons, we hypothesized ubiquitination may impair VASP tetramerization and, subsequently, VASP activity. We performed mass photometry to quantify the molecular mass of VASP. As a monomer of TAMRA-VASP is 41.3 kDa, the estimated tetrameric mass is 123.9 kDa. We detected a single peak corresponding to a tetramer for unmodified VASP centered around 141 kDa. A similar single population was observed for mUb-K240 VASP, mUb-K286 VASP, and mmUb-K240/K286 VASP (Fig 1E). No significant peak shifts were observed beyond the margin of error. Thus, mUb-K240, mUb-K286, and mmUb-K240/K286 VASP each maintain tetramerization.

### mmUb-K240/K286 VASP exhibits decreased actin filament binding and bundling

With purified, ubiquitinated VASP tetramers in hand, we began testing the ability of VASP-Ub to bind actin filaments through high speed co-sedimentation assays (Fig 2A). At this centrifugation speed, actin filaments—and any filamentous binding partners—are pelleted. Consistent with previous results (Reinhard et al., 1992; Breitsprecher et al., 2011), the majority of unmodified VASP co-sedimented with actin filaments (Fig 2B,C). A single ubiquitination modification at K286 (mUb-K286) did not significantly change the amount of VASP present in the pellet, however, a small but significant decrease was observed with 600 nM of mUb-240. Furthermore, there was a significant decrease in the amount of mmUb-K240/K286 that co-sedimented with actin at 400 nM and 600nM (Fig 2C). This suggests that VASP multi-monoubiquitination decreased filamentous actin binding affinity.

**Fig. 2.**
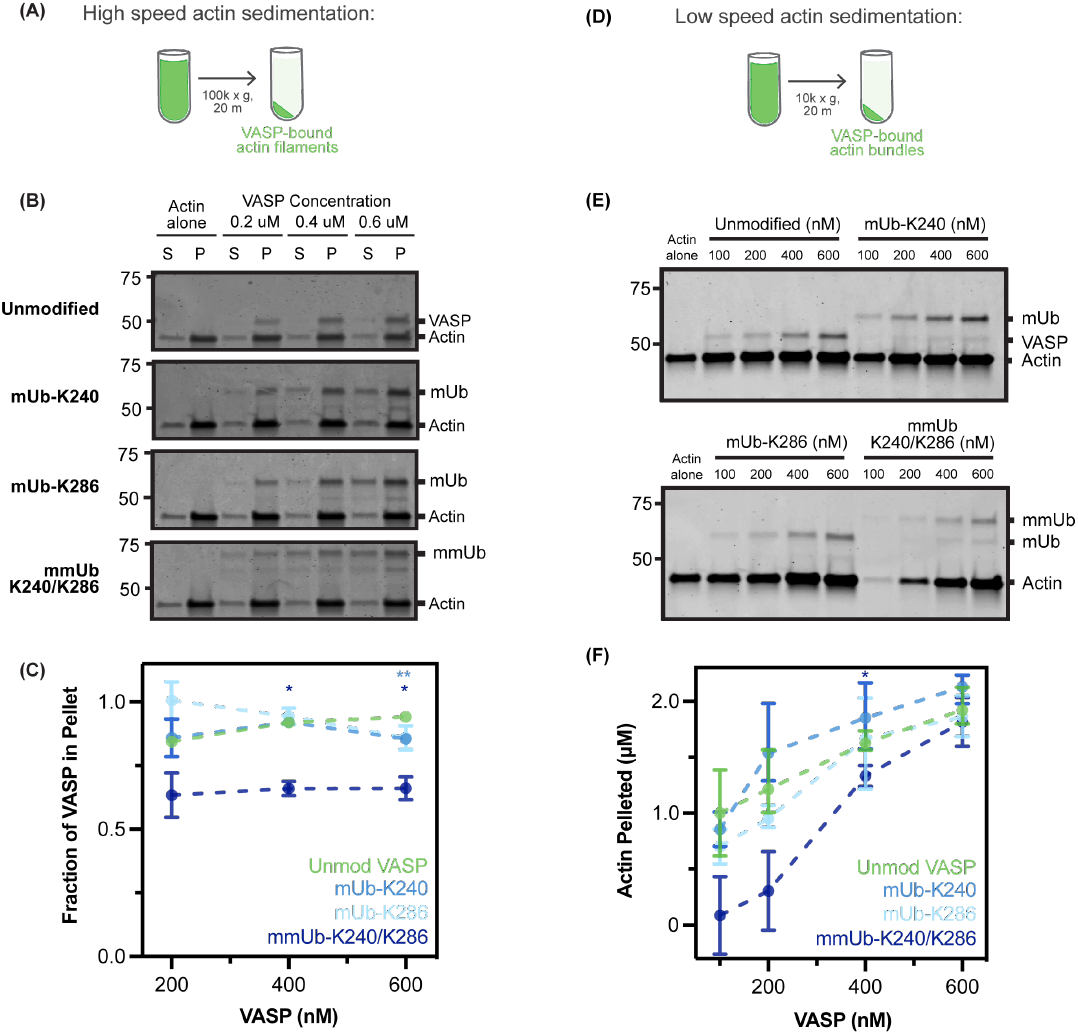
Multi-monoubiquitination of VASP impairs actin bundling and binding. (A) High-speed actin co-sedimentation assay utilizing 1 μM actin and varying concentrations of VASP (B) Coomassie gels of supernatant and pellet fractions following centrifugation. (C) Quantification of VASP localized to the pellet fraction, determined by densitometry of Coomassie gels. Data points represent the mean ± SEM from 3 independent experiments. Statistics were calculated with a two-way ANOVA with a Geisser-Greenhouse correction and the Dunnett’s multiple comparisons test. Unmodified VASP v. mUb-K240/K286: 400 nM = 0.0315, 600 nM = 0.0466. Unmodified VASP v. mUb-K240: 600 nM = 0.0089. (D) Low-speed actin co-sedimentation assay utilizing 2 μM actin and varying concentrations of VASP. (E) Coomassie gels of the pellet fraction following centrifugation. (F) Quantification of actin localized to the pellet fraction, determined by densitometry of Coomassie gels. Data points represent the mean ± SEM from 3 independent experiments. Statistics were calculated with a mixed effects model with a Geisser-Greenhouse correction and the Dunnett’s multiple comparisons test. Unmodified VASP v. mUb-K240/K286: 200 nM = 0.06399, 400 nM = 0.0399.

VASP also promotes actin filament bundling. We completed similar assays at low centrifugation speeds to evaluate the ability of VASP to promote bundling of actin filaments (Fig 2D). Compared to actin alone, unmodified VASP increased bundled actin in a concentration dependent manner (Fig 2E, F). We observed similar actin bundling activity in the presence of mUb-K240 and mUb-K286. A decrease in actin bundling activity by mmUb-K240/K286 at lower concentrations was apparent, although this change was only significant at 400 nM. By 600nM—a concentration at which the actin bundling activity of unmodified VASP was saturated—the bundling activity of mmUb-K240/K286 recovered (Fig 2E, F).

### mmUb-K240/K286 VASP exhibits impaired ability to accelerate actin filament elongation

The acceleration of actin filament elongation by VASP requires two separate actin binding events. First, VASP must bind to the growing “barbed” end of the actin filament through the FAB (filamentous actin binding domain). Secondly, VASP binds monomeric actin through the GAB (G-actin binding domain). This presence of four GAB domains—each able to bind an actin monomer—is thought to increase the local pool of monomeric actin available for polymerization at the barbed end, accelerating elongation (Hansen and Mullins, 2010). Due to the proximity of the identified ubiquitination sites to the GAB (K240-Ub) and the FAB (K286-Ub), we hypothesized that ubiquitinated VASP may also exhibit a deficit in filament elongation. We used seeded pyrene assays—in which labeled actin monomers were added to pre-formed actin filaments—to specifically examine the elongation step of actin polymerization (Fig 3A). We analyzed the first 300 s of the assay to determine the initial elongation rate at range of VASP concentrations (Fig 3B). Consistent with previous studies (Harker et al., 2019), unmodified VASP displayed a concentration-dependent increase in the rate of actin elongation before reaching a plateau.

**Fig. 3.**
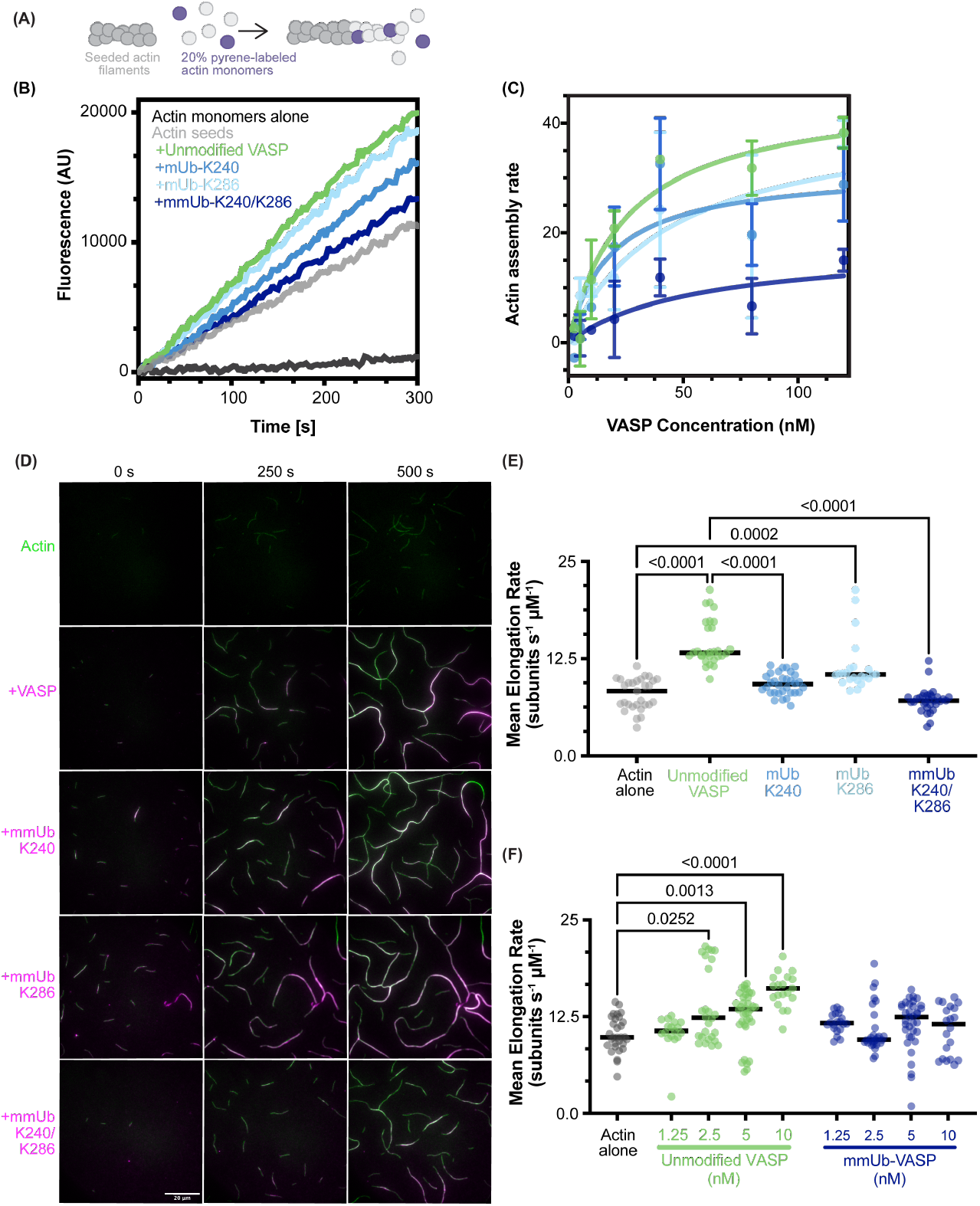
Mono- and multimono-ubiquitination of VASP impairs elongation of actin filaments. (A) Schematic of pyrene actin assays. (B) Fluorescent traces of 0.5 μM (20% pyrene-labeled) actin monomer elongation from preformed actin seeds (5 μM) in the presence of 80 nM VASP. (C) The initial elongation rate from seeded pyrene elongation assays (calculated from the first 300 s of the assay) plotted against VASP concentration. A one-site binding curve was fit to each data set. The elongation rate of actin seeds alone was subtracted from each data point. Each data point contains 1-4 measurements collected across 3 independent experiments. (D) Polymerization of 1.5 μM actin (10% AlexaFluor-488 labeled) visualized with TIRF microscopy for ten minutes in the presence of 25 nM VASP. (E) Quantification of actin filament elongation rates from the timelapse images. N = 23-32 filaments per condition from 2-3 experiments. Lines represent median elongation rate. Elongation rate was calculated assuming 375 subunits were added per μm. Statistical significance calculated with the Kruskal-Wallis test and Dunn’s multiple comparison test. (F) Quantification of actin filament elongation rates at a range of VASP and mUb240/286 concentrations. N = 17-36 filaments from 2-4 experiments for each data point. Lines represent median elongation rate. Indicated p values were calculated with the Kruskal-Wallis test and Dunn’s multiple comparison test.

mUb-K240 or mUb-K286 did not significantly change the plateaued maximal actin elongation rate. We also estimated similar K_D_ values as unmodified VASP for both modifications (Table 1), suggesting the affinity of VASP for the barbed end of the actin filament was not detectably altered by either monoubiquitination event. Strikingly, the cumulative effect of both monoubiquitins in mmUb-K240/K286 severely reduced the rate of actin elongation, with the values remaining close to actin alone. Although actin elongation was impaired by multimonoubiquitination, results from this bulk assay over multiple concentrations of mmUb-K240/K286 also suggested a similar K_D_ value as unmodified VASP (Table 1). However, these calculations are limited by the uncertainty of the maximum elongation rate for this protein.

To examine changes in actin polymerase activity in greater detail, we utilized total internal reflection fluorescence microscopy (TIRFM) to determine the elongation rate of individual actin filaments. Each VASP construct (25 nM) was mixed with 1.5 μM actin monomers (10% AF-488 labeled) and spontaneous actin assembly was imaged for ten minutes (Fig 3D, Supplemental Movie 1). Focusing on the elongation rates of single actin filaments and parallel actin bundles, quantification was largely consistent with bulk pyrene-actin polymerization. Unmodified VASP increased the mean actin elongation rate 1.8-fold compared to actin alone. Although the activity of mUb-K286 was similar to unmodified VASP, both mUb-K240 and mmUb-K240/K286 exhibited a decreased rate of elongation (Fig 3D, E). Although we did not detect a significantly changed elongation rate for mUb-K240 by pyrene actin elongation assays, we account this disparity to the higher sensitivity of the TIRF assays.

As the time-lapse movies progressed, we observed VASP-mediated bundling of actin filaments. In the presence of unmodified VASP, actin bundles quickly predominated the field of view, obscuring the polymerization of single actin filaments (Fig 3D, 250 and 500 s). Consistent with the low speed co-sedimentation assays, the impaired ability of mmUb-K240/K286 to bundle actin filaments was visibly apparent at 500 s.

To further probe the striking difference in actin polymerase activity between unmodified and mmUb-K240/K286 VASP, we measured the actin elongation rate in the presence of either protein at a range of concentrations (1.25 nM-10 nM). Although unmodified VASP exhibited a clear concentration-dependent increase in the elongation rate, mmUb-K240/K286 failed to significantly increase the rate of elongation compared to actin alone (Fig 3E, Supplemental Movie 2). Collectively, both seeded pyrene assays and TIRF data support the conclusion that the enhancement of new actin monomer addition is impaired by multi-monoubiquitination of VASP.

### Ubiquitinated VASP maintains anti-capping activity

Capping protein binds to the barbed ends of actin filaments and terminates actin polymerization (Cooper and Pollard, 1985). Previous work demonstrated that VASP competes with capping protein for binding and thus protects the barbed end (Barzik et al., 2005). The anti-capping activity of Ena/VASP proteins is essential for their unique contribution to maintaining the actin cytoskeleton. In cultured cells, loss of VASP was associated with short, highly branched actin filaments (Bear et al., 2002). Previous work demonstrated that both GAB and FAB domain function is required for anti-capping activity (Barzik et al., 2005), suggesting that ubiquitination at either K240 or K286 may modulate the anti-capping activity of VASP.

To evaluate the anti-capping activity of VASP-Ub, we performed seeded pyrene assays in the presence of 5 nM mammalian capping protein (Fig 4A). Compared to actin seeds alone, the addition of capping protein severely reduced actin elongation (Fig 4B). As reported previously (Bear et al., 2002; Barzik et al., 2005), the addition of VASP counteracted capping protein in a concentration dependent manner. All three ubiquitinated versions of VASP enhanced actin elongation over capping protein and actin (Fig 4B, C), suggesting that mono- and multi-monoubiquitinated VASP maintained anti-capping activity. Like pyrene assays in the absence of capping protein, our curve fitting calculated similar K_D_ values, further supporting a similar affinity of unmodified and ubiquitinated VASP for the barbed end (Table 2).

**Fig. 4.**
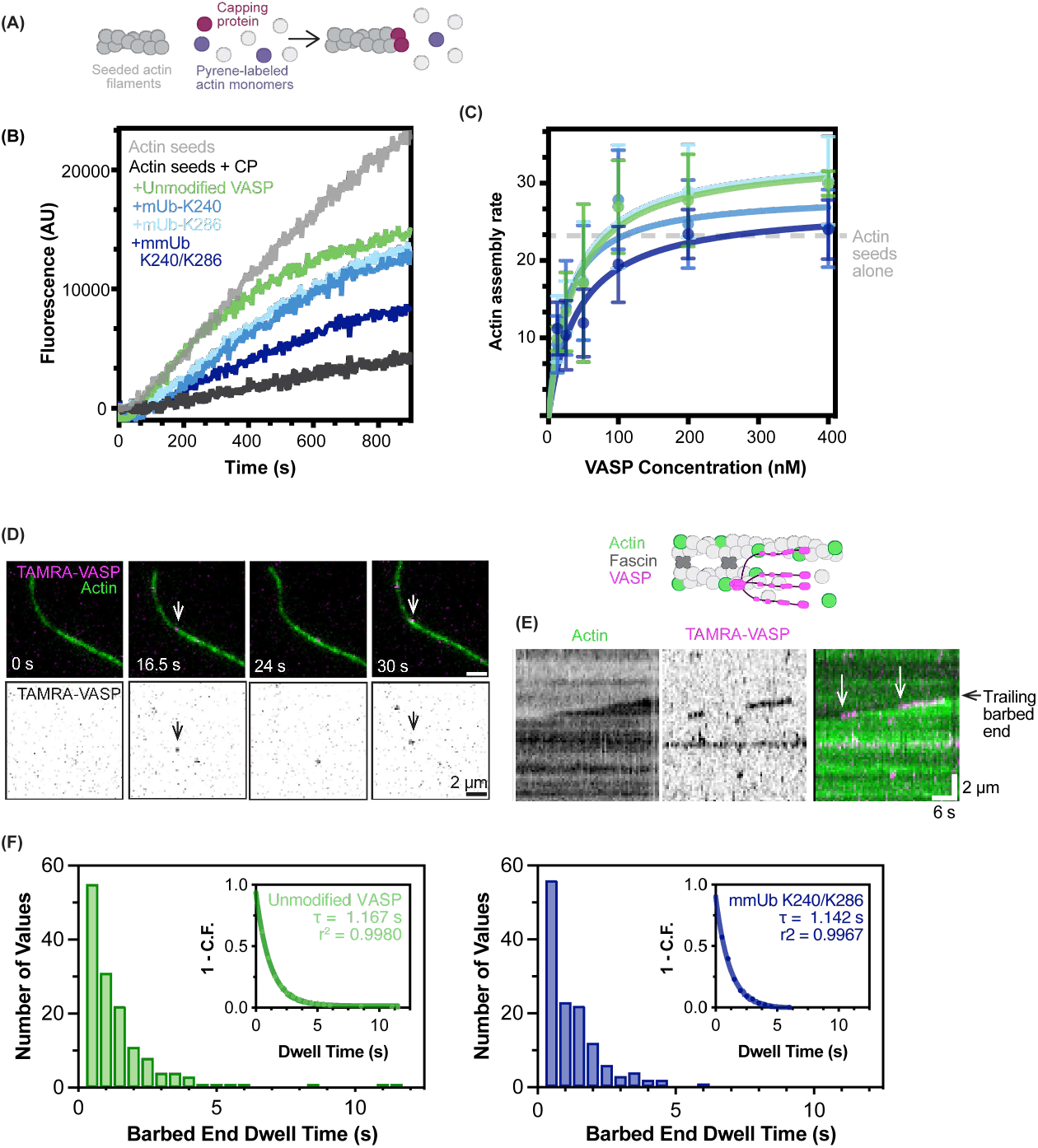
Multi-monoubiquitination of VASP does not change barbed end binding or anti-capping activity. (A) Schematic of pyrene actin assays in the presence of capping protein (B) Fluorescent traces of 0.5 μM (20% pyrene-labeled) actin monomer elongation from preformed actin seeds (5 μM) in the presence of 5 nM capping protein (CP) and 50 nM VASP. (C) The initial elongation rate from seeded pyrene elongation assays (calculated from the first 300 s of the assay) plotted against VASP concentration. A one-site binding curve was fit to each data set. The dotted line represents the average elongation rate of actin seeds alone. Each data point contains 3-5 measurements collected across 4 independent experiments. (D) TIRFM demonstrating localization of 1 nM TAMRA-VASP to the barbed end of a trailing actin filament (1.5 μM actin, 10% AlexaFluor-488 labeled) in a fascin-mediated actin bundle. Arrowheads indicate VASP bound to the trailing barbed end. (E) The growth of the trailing actin filament and the dynamic localization of VASP to the barbed end in (D) was visualized with kymograph analysis. Arrowheads denote two processive VASP binding events at the barbed end of 2-filament bundles. These highly processive binding events are rare, with the majority of events lasting < 1 s. (F) Frequency distribution of VASP barbed end binding events. Graph inset demonstrates (1-Cumulative Frequency) vs. dwell time fit with a one-phase decay curve. N = 145 events (unmodified VASP) and 131 events (mUb-K240/K286) from 2 experiments.

### mmUb-K240/K286 VASP maintains association with barbed end

We observed striking deficits in the ability of mmUb-K240/K286 to increase the elongation rate of actin filaments, yet the protein still possessed anti-capping activity. To determine if VASP mmUb-K240/K286 maintained interaction with the barbed end of actin filaments, we quantified the lifetime of VASP puncta bound to the barbed end in single molecule TIRFM imaging. This experiment is complicated by the relatively short lifetime of human VASP, previously measured at τ_1_ = 1.45 s (Hansen and Mullins, 2010) and τ_1_ = 1 s (Harker et al., 2019). However, the lifetime of VASP is increased on the trailing filament ends of fascin-bundled actin (τ_1_ = 2.6 s and 4.2 s for bundles containing two and three or more filaments, respectively) (Harker et al., 2019). Due to the maximum acquisition speed of the microscope (0.5 s/ frame), we utilized fascin-bundled actin to more accurately quantify the lifetimes of unmodified VASP and VASP mmUb-K240/K286 on the trailing filament of a two-filament bundle (Fig 4D, E). We estimated the lifetime of unmodified VASP at τ_1_ = 1.167 s (95 % CI = 1.094 to 1.245 s). We observed a similar dwell time (τ_1_ = 1.142 s (95 % CI = 1.007 to 1.301 s) of mmUb-K240/K286 at the barbed end (Fig 4F). As inverse of puncta lifetime is a measurement of the k_off_ rate, we infer that multimonoubiquitination did not affect the k_off_ rate of VASP.

### mmUb-K240/K286 VASP exhibits impaired function and localization in cells

Previously, we demonstrated that the mutation of lysine residues in VASP—reducing VASP ubiquitination—resulted in increased filopodial number and stability in cultured murine neurons (Menon et al., 2015; Boyer et al., 2020). However, there is no primary coding sequence mutation that can mimic the ubiquitination of VASP in cells. To examine how ubiquitination of VASP altered its function and localization in cells, we electroporated purified, ubiquitinated VASP into cultured cells. Since Ena/VASP proteins can hetero-tetramerize (Riquelme et al., 2015), we utilized MV^D7^ cells (Bear et al., 2000), a mouse embryonic fibrob-last line lacking all Ena/VASP family members, to simplify experiment interpretation. In this line, Enah (the gene encoding Mena) and Vasp were genetically deleted before subsequent selection for undetectable levels of EVL protein. Following electroporation of TAMRA-labeled VASP, cells were permitted to spread for thirty minutes before fixation.

Previous work characterized spreading of MV^D7^ cells into three phenotypes: smooth-edged cells containing a clear lamellipodia; ruffled cells characterized by actin ruffles; and filopodial-rich cells (Applewhite et al., 2007). (Fig 5A, Fig S2A). Using a low concentration of TAMRA-VASP electroporated into cells, we observed no changes in the distribution of these three classes from cells that were electroporated with buffer alone. However, electroporation of mmUb-K240/K286 VASP decreased the proportion of filopodial-like cells and subsequently increased the proportion of ruffled cells (Fig 5C). We observed no significant changes in filopodia density (in cells classified as filopodial), cell area, or phalloidin staining following electroporation of either construct (Fig S2B).

**Fig. 5.**
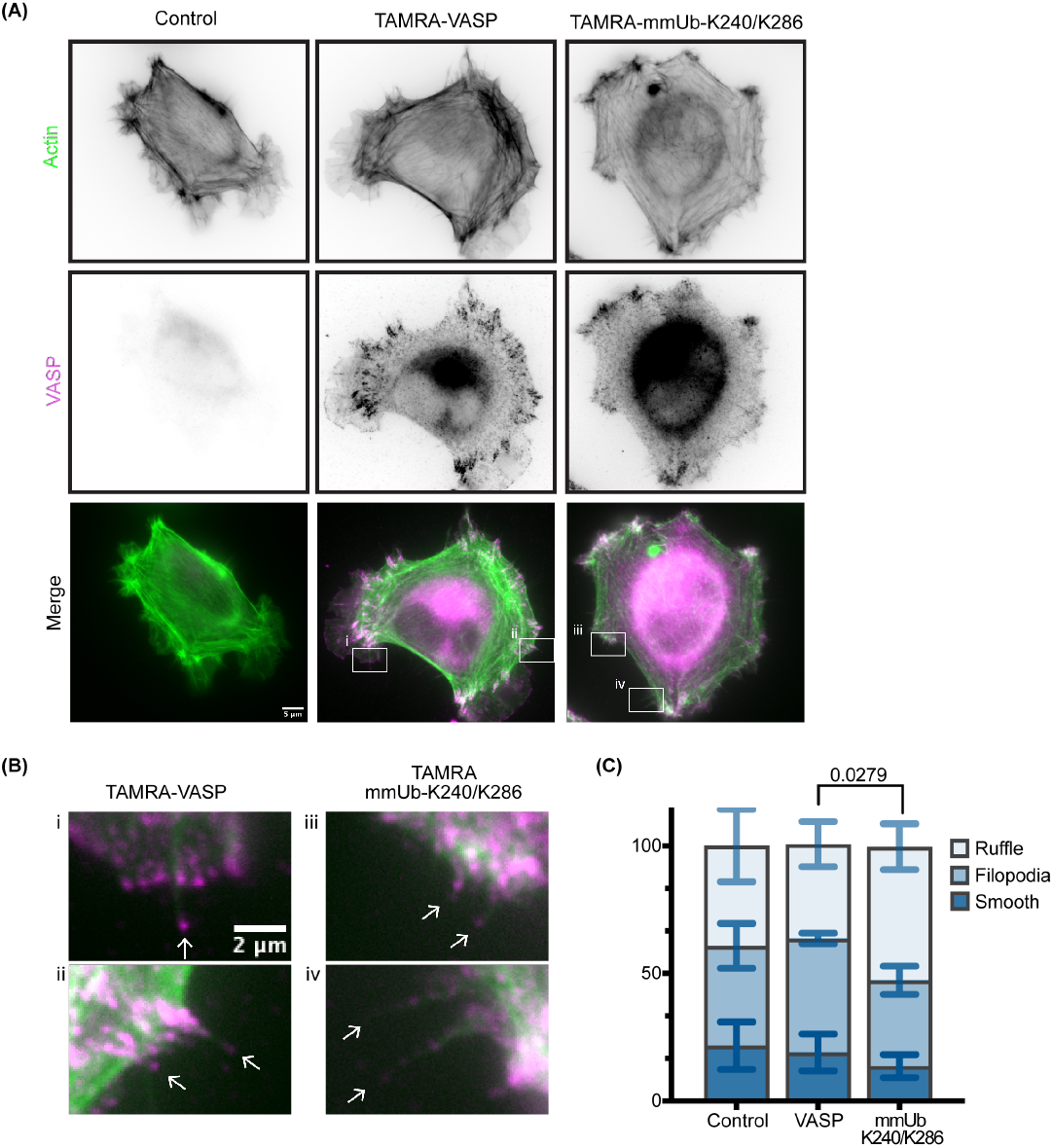
Multi-monoubiquitination of VASP enhances actin ruffling during cell spreading. (A) Representative widefield images of fixed MVD7 cells with filopodial spreading morphology. Cells were electroporated with TAMRA-VASP (or buffer control) and allowed to spread for 30 minutes on fibronectin. (B) Insets from (A) highlighting VASP localization to the tips of filopodia. (C) Classification of cell spreading into smooth-edged, filopodial, or ruffled phenotypes. The plot indicates mean percentage ± SD of each classification across 3 experiments. A Chi-square test for the goodness of fit was used to compare outcomes with a Bonferroni correction. 92-104 cells were classified across the experiments.

We observed localization of both TAMRA-VASP and TAMRA-mmUb-K240/K286 to filopodial tips, lamellipodia, and focal adhesions in MVD7 cells (Fig 5B). To further examine this localization, we electroporated each protein into MV^D7^ cells stably expressing GFP-VASP (Fig 6A). We observed clear co-localization of TAMRA-VASP and GFP-VASP at all three of these actin-based structures. Although TAMRA-mmUb-K240/K286 was present as well, its enrichment at each of these structures appeared dimin-ished. Quantification of raw values indicated significantly less mmUb-K240/K286 VASP localized to filopodia tips compared to TAMRA-VASP (Fig 6B).

**Fig. 6.**
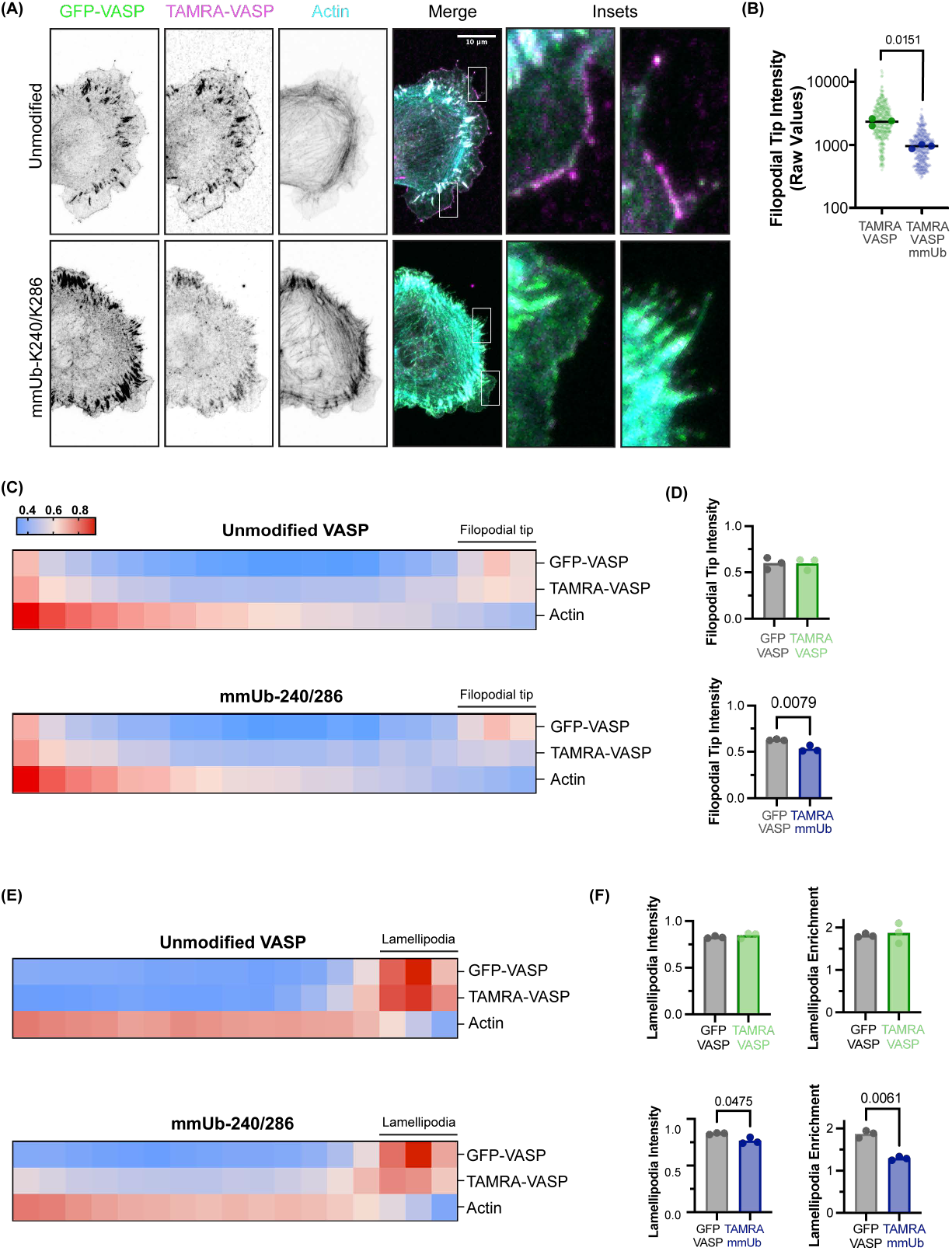
Multi-monoubiquitination of VASP reduces enrichment at both filopodial tips and lamellipodia. (A) Representative maximum projection images of fixed MVD7 cells stably expressing GFP-VASP and electroporated with TAMRA-VASP. (B) Raw intensity values of average filopodial tip intensity from linescans. The filopodial tip was defined as the first four pixels (0.255 μm) in a linescan drawn from tip to base. Each large data point represents the average from one independent experiment with smaller data points representing individual filopodia. The line represents the mean of three independent experiments. Statistics were calculated with a paired t-test. N = 34 (Unmodified) and 35 (mmUb-K240/K286 VASP) cells across 3 experiments for all filopodia measurements. (C) Heatmap localization of proteins from throughout the length of the filopodia. The fluorescence intensity of each protein with a filopodium was normalized and the length binned. (D) Quantification of normalized VASP intensity at the filopodial tip (first three bins). Each data point represents the mean of one independent experiment and the bar graph extends to the mean of three experiments. Statistics were calculated with a paired t-test. (E) Heatmap localization of proteins at the lamellipodia. Line scans were drawn from the cell periphery and extended approximately 1.5 μm into the cell. The fluorescence intensity of each linescan was normalized and the length binned. (F) Quantification of normalized VASP intensity at the lamellipodia (first three bins). Each data point represents the mean of one independent experiment and the bar graph extends to the mean of three experiments. Lamellipodia enrichment was calculated as the intensity at the first three bins normalized to the intensity and the subsequent fourteen bins. Statistics were calculated with a paired t-test. N = 31 (Unmodified) and 33 (mmUb-K240/K286 VASP) cells across 3 experiments for all lamellipodia measurements.

To compare the distribution of unmodified VASP and mmUb-K240/K286 VASP that did localize within filopodia, we normalized VASP and actin levels along the length of filopodia. As previously reported (Applewhite et al., 2007; Rottner et al., 1999; Lebrand et al., 2004), GFP-VASP was enriched at the tips of filopodia. In contrast, phalloidin staining was most intense at the base of the filopodia, tapering in intensity towards the tip. Whereas TAMRA-VASP mirrored GFP-VASP localization, the filopodial tip intensity of TAMRA-mmUb-K240/K286 was diminished (Fig 6C,D). Similarly, linescans were utilized to evaluate VASP localization within the lamellipodia. Both GFP-VASP and TAMRA-VASP strongly colocalized at the cell periphery. Although TAMRA-mmUb-K240/K286 was still present at the lamellipodia, the fluorescence intensity was decreased relative to GFP-VASP. Furthermore, the ratio of fluorescence intensity at the lamellipodia to the adjacent medial lamella was also decreased (Fig 6E,F). Thus, multi-monoubiquitination of VASP reduced protein enrichment at both the filopodial tip and lamellipodia.

## Conclusions

The actin cytoskeleton performs many essential cellular functions; thus actin polymerization must be tightly regulated. Due to their rapid occurrence and reversible nature, post-translational modifications are ideal candidates to help modulate actin dynamics yet, with the exception of phosphorylation, are relatively understudied. Previously published work demonstrated non-degradative ubiquitination of VASP reduced filopodial number and stability (Menon et al., 2015; Boyer et al., 2020). Although the mechanistic impact of VASP ubiquitination was unclear, cellular assays suggested VASP was mono or multi-monoubiquitinated (Boyer et al., 2020). Here we identified prominent ubiquitination of VASP at K240 and K286, and generated recombinant forms of ubiquitinated VASP to interrogate actin regulatory functions. Whereas mUb-K286 did not significantly affect VASP function in any of our tested assays, monoubiquitination of mUb-K240 significantly reduced the ability of VASP to elongate actin filaments. A multi-monoubiquitinated construct (mmUb-K240/K286) showed significant decreases in its ability to bind, bundle, and elongate actin filaments and localize to filopodia tips and lamellipodia.

### Mechanism of impairment of polymerase activity

VASP enhances actin filament elongation, reflecting its ability to bind both G-actin and F-actin through respective GAB and FAB domains (Hansen and Mullins, 2010). We observed decreases in the ability of both mUb-K240 and mmUb-K240/K286 to accelerate elongation of actin filaments. Seeded pyrene assays suggested all three ubiquitinated VASP proteins similarly associated with the barbed end of the actin filament, with a similar K_D_ calculated for each protein. However, this interpretation is limited by the sensitivity of the pyrene assay. The 95% confidence interval for all three ubiquitinated proteins spanned a large range and we were unable to calculate an upper limit for mmUb-K240/K286. In contrast, we observed more dramatic differences in the elongation rates calculated through more sensitive single molecule TIRFM assays. At the range of concentrations tested (1.25-25 nM), we did not observe a significantly different elongation rate for mmUb-K240/K286 compared to actin alone. Although this could indicate that the binding affinity of mmUb-K240/K286 for the barbed end was decreased, several lines of evidence indicate that the protein maintains a reasonable association with the barbed end. First, mmUb-K240/K286, along with both single mUb VASP, maintained anti-capping activity, indicating interaction with barbed end. Second, we calculated similar lifetimes of unmodified VASP and mmUb-K240/K286 bound to the barbed end. Although our measurements were limited by the speed of microscopy acquisition, we did not detect significant differences between the two proteins. Collectively, pyrene elongation assays (both in the presence and absence of capping protein) and single molecule TIRFM assays suggest similar barbed end binding dynamics for unmodified VASP and mmUb-K240/K286. As such, we postulate other mechanisms by which mmUb-K240/K286 may impair the acceleration of actin filament elongation. First, this deficit may arise from an impaired ability of ubiquitinated VASP to interact with monomeric actin. Of note, K240 is adjacent to the GAB and a crystal structure of this region demonstrates L234 is an essential residue for monomeric actin binding (Ferron et al., 2007). Although this structure suggests K240 lies in an unstructured region that does not directly bind the actin monomer, ubiquitin is 8 kD and has the potential to impair the binding between L234 and an actin monomer through steric hindrance. However, previous work indicates actin monomer binding is required for effective barbed end localization (Hansen and Mullins, 2010) and we did not detect changes in barbed end dwell time.

Alternatively, it is possible that the GAB maintains its monomeric actin binding activity and mUb-K240 may sterically impede the transfer of monomeric actin onto the filament. As filament elongation is severely reduced with the addition of a second ubiquitination site at K286, this supports a model with impaired transfer of the actin monomer onto the filament. Interestingly, pyrene assays in the presence of 5 nM capping protein suggest that mmUb-K240/K286 maintains anti-capping activity. In contrast, VASP constructs containing either GAB mutations (R232E,K233E) or FAB mutations (L}256-273) completely lost the ability to protect barbed ends from capping protein (Barzik et al., 2005), suggesting a functional GAB and FAB are needed for anti-capping activity of VASP. As such, these results support that mmUb-K240/K286 maintains both GAB and FAB binding and instead there is a deficit in the transfer of actin monomers onto the filament.

N-terminal to the GAB domain, VASP also contains a poly-proline region that binds profilin-actin. Previously published in vitro experiments have made mixed conclusions on the utilization of profilin-actin by Ena/VASP proteins due to differences in experimental design (Hansen and Mullins, 2010; Breitsprecher et al., 2008; Brühmann et al., 2017). However, a recent cellular study suggests that Ena/VASP proteins do require profilin-actin for leading edge actin polymerization (Skruber et al., 2020). Although ubiquitination at K240 or K286 is unlikely to impair VASP-profilin binding within the polyproline region, whether ubiquitination may block the transfer of profilin-actin to the GAB and FAB is not known.

### Multi-monoubiquitination of VASP regulates cell dynamics

Precise regulation of the actin cytoskeleton is required for cell morphology and motility changes. However, cells must quickly adapt to a variety of stimuli, including extracellular cues. In this work, we electroporated TAMRA-VASP and TAMRA-mmUb-K240/K286 VASP into MVD7 cells. In the presence of TAMRA-VASP mmUb-K240/K286, we observed changes in cell spreading morphology and intracellular localization of VASP. Because VASP and TRIM9 colocalize at filopodia tips, we propose VASP monoubiquitination is mechanism to rapidly control filopodial activity. Furthermore, this modification is reversible, avoiding the lengthy cycles of protein synthesis/degradation. E3 ligases exist in a balance with deubiquitinase (DUB) enzymes. Although we have demonstrated pharmacologically blocking DUB activity prevents VASP deubiquitination (Menon et al., 2015), the DUB for VASP is unknown. Identification of this enzyme is crucial to fully comprehend the cycle of VASP regulation in neurons, and in turn, cytoskeletal regulation.

Previously published results demonstrated the E3 ubiquitin ligase TRIM9 is required for ubiquitination of VASP in cultured neurons and HEK293 cells (Menon et al., 2015; Boyer et al., 2020). However, in the absence of TRIM9, residual VASP ubiquitination remained, suggesting redundancy with other E3 ligases (Boyer et al., 2020). The discovery of additional TRIM9 substrates in developing neurons remains an open area of inquiry. Previously, we demonstrated the netrin-1 receptor DCC is ubiquitinated in a TRIM9-dependent manner (Plooster et al., 2017). Although the type of ubiquitin modification is still unknown, molecular weight shifts and protein stability similarly suggested that it is not degradative polyubiquitination. As DCC also localizes to the tips of filopodia (Shekarabi and Kennedy, 2002), this evokes a multi-faceted mechanism by TRIM9 to regulate filopodia function in neurons. A recent BioID experiment also identified a multitude of potential TRIM9 binding partners (Menon et al., 2021, 2020), yet additional work is required to validate these candidates as TRIM9 ubiquitination substrates.

TRIM9 has been studied primarily in the context of neuronal development and neurodegeneration (Menon et al., 2015; Boyer et al., 2020; Plooster et al., 2017; Tanji et al., 2010; Winkle et al., 2016; Zeng et al., 2019; Winkle et al., 2016; Do et al., 2018). However, TRIM9 is also expressed in macrophages (Tokarz et al., 2017; Carthagena et al., 2009) and various forms of cancer (Yang et al., 2020a; Lin et al., 2023; Zhang et al., 2023; Mishima et al., 2015). Although the alteration of TRIM9 protein levels has been shown to affect cell proliferation, motility, and inflammation in cell lines, substrates of TRIM9 have not been identified in these studies. Of note, filopodia play a critical role in cell migration in various cancers (Jacquemet et al., 2015) and contribute to phagocytosis in macrophages (Horsthemke et al., 2017). However, whether TRIM9 localizes to filopodia in these cells and is capable of ubiquitinating VASP is unknown.

Non-degradative ubiquitination also modulates several other cytoskeletal pathways. For example, two isoforms of the oncogene Ras, a small GTPase that regulates numerous cytoskeleton effectors, are monoubiquitinated. Monoubiquitination of H-Ras at K117 increased GTP-GDP exchange, leading to increased Ras activity (Baker et al., 2013). Likewise, monoubiquitination of K-Ras at K147 increased K-Ras activation by blocking binding to GTPase activating (GAP) proteins (Baker et al., 2013; Sasaki et al., 2011). In contrast, mono-ubiquitination of K-Ras at K104 did not impair biochemical activity of the protein and the function of this ubiquitination site is still unknown (Yin et al., 2020). Furthermore, the actin bundling protein fascin is monoubiquitinated by the E3 ligase Smurf1 at K247 and K250. Biochemical studies showed mUb-K250 decreased actin bundle thickness and bundle persistence, while transfection of non-ubiquitinatable fascin increased cell migration in an adenocarcinoma cell line (Lin et al., 2016).

### Deciphering the ubiquitination reaction

Ubiquitination is a complex cascade, requiring the sequential activity of an E1 ubiquitin-activating enzyme, an E2 conjugating enzyme, and an E3 ubiquitin ligase. As an E3 ligase, TRIM9 catalyzes the transfer of ubiquitin from the E2 enzyme onto the substrate. E3 ligases add specificity to ubiquitination reactions; more than 600 mammalian E3 ligases have been identified (Yang et al., 2021). However, the activity of an E3 ligase can change based on which E2 conjugating enzyme it partners with. Previous in vitro biochemical experiments have identified E2 partners of TRIM9—including UbcH5b/UBE2D2 and UBE2G2 (Tanji et al., 2010; Napolitano et al., 2011)—however, how E2 partnership informs TRIM9 substrate selectivity in vivo is unknown.

Likewise, how TRIM9 activity is regulated is unknown—whether by post-translational modifications or additional protein-protein interactions. E3 ubiquitin ligases are well-appreciated to form larger complexes to efficiently and specifically ubiquitinate their targets. For example, Melanoma Antigen Gene (MAGE) proteins bind certain E3 ligases and regulate E3 activity through a variety of mechanisms, including E2 protein stabilization and substrate binding (Lee and Potts, 2017; Yang et al., 2020b).

### Ubiquitination and phosphorylation crosstalk

Post-translational modifications are essential regulators of protein stability, activity, and localization. Unlike ubiquitination, the mechanistic impact of VASP phosphorylation is well studied in numerous cell types. For example, phosphorylation of VASP at S239 and T278 by cGMP-dependent protein kinase (PKG) reduced actin polymerization in vitro and filamentous actin levels in HEK293 cells (Benz et al., 2009). Furthermore, work in human colon cancer carcinoma cell lines suggested pS239 reduces VASP enrichment at filopodia tips and decreases filopodia length (Zuzga et al., 2012). Of note, cross talk may also occur between post-translational modifications, including between ubiquitination and phosphorylation. For example, phosphorylation of the small GTPase RhoA at Ser188 by PKA/PKG blocks ubiquitination and proteasomal degradation of the protein (Rolli-Derkinderen et al., 2005). In contrast, phosphorylation of RhoA by the kinase Erk2 promotes RhoA ubiquitination (Wei et al., 2013). Furthermore, our previous work suggests TRIM9-mediated ubiquitination of DCC reduces phosphorylation at Tyr1418 (Plooster et al., 2017).

With these results in mind, we note that the VASP ubiquitination site (K240) is located directly next to a phosphorylation site (S239) and both modifications are associated with negatively regulating VASP function. As such, we ponder this biological redundancy. However, we can speculate that that VASP mUb-K240 is a more specific signaling pathway compared to S239-P. The kinase PKG has a prolific number of substrates and is stimulated by cGMP, a messenger with effectors throughout the body. Indeed VASP is phosphorylated in this pathway, but it certainly will not be the only protein to be modified. In contrast, Trim9 is predominantly expressed in neurons and glia. Furthermore, TRIM9 shows specific enrichment at the filopodial tip, providing precise local access to VASP. We previously reported VASP is deubiquitinated in the presence of the guidance cue, netrin (Menon et al., 2015). As mentioned above, the netrin receptor DCC also localizes to the tips of filopodia (Shekarabi and Kennedy, 2002) and is ubiquitinated in TRIM9-dependent manner (Plooster et al., 2017). Collectively, TRIM9-mediated ubiquitination may serve as a filopodial-specific negative regulator of cytoskeletal remodeling.

## Data Availability

Proteomics data is deposited in PRIDE (https://www.ebi.ac.uk/pride/) and can be accessed with the following information:

Project Name: Ubiquitination of the actin polymerase VASP Project accession: PXD041196

Reviewer account details:

Username

Password: 2i9n20l4

### Experimental Model and Subject Details

HEK293 cells were obtained from Simon Rothenfußer (Klinikum der Universitat München, München, Germany) as previously described (Menon et al., 2015). Cells were cultured in DMEM (Gibco Cat #11965092) with 10% FBS (Hyclone) and maintained at 5% CO2/37°C. MV^D7^ cells (Bear et al., 2000) were a gift from Frank Gertler (MIT). Cells were maintained in DMEM supplemented with GlutaMAX (ThermoFisher Cat #35050061), 15% FBS (ThermoFisher Cat #FB12999102), and 50 U/mL of interferon-γ (Sigma Cat #IF005). Cells were cultured at 32°C with 5% CO2.

## Methods

### Plasmid transfections

HEK293 cells were transfected with Lipofectamine200 (ThermoFisher Scientific) according to the manufacturer’s instructions.

### Plasmids, antibodies, and reagents

Creation of Myc-VASP was described previously (Menon et al., 2015; Boyer et al., 2020). pRK5-HA-Ubiquitin-WT was a gift from Ted Dawson (Addgene plasmid 17608; http://n2t.net/addgene:17608; RRID:Addgene_17608) (Lim et al., 2005). pGEXTEV Flag PS-SSSS-UBG75C creation was previously described (Brown et al., 2016).

Cysteine-light His-VASP (His-VASPCCC-SSA) containing an N-terminal KCK motif was a gift from Dyche Mullins (UCSF) (Hansen and Mullins, 2010). To facilitate sortase labeling, the KCK motif was mutated to a GGG motif using Quik-Change mutagenesis. VASP K240C and VASP K286C were also generated using Quik-Change mutagenesis. VASP K240C, K286C was created by Azenta Life Sciences through site-directed mutagenesis.

Primary antibodies used were a mouse monoclonal against c-Myc (9E10) (1:2000 WB, Hybridoma serum purified in-house) and a rabbit polyclonal against the HA-tag (1:250 WB, ThermoFisher Scientific Cat 71-5500). Fluorescent secondary antibodies included IRDye 680LT Goat anti-Rabbit IgG Secondary Antibody (1:10,000 WB, LI-COR Cat NC9030093) and IRDye 800CW Goat anti-Mouse IgG Secondary Antibody (1:10,000 WB, LI-COR Cat NC9401841).

Other reagents included PR-619 (Fisher Scientific Cat #66-214-125), MG132 (American Peptide Company #81-5-15), Alexa Fluor™ 488 Phalloidin (ThermoFisher Scientific Cat #22323), Alexa Fluor™ 647 Phalloidin (ThermoFisher Scientific), bismaleimidoethane (ThermoFisher Scientific Cat #22323), and paraformaldehyde (ThermoFisher Scientific Cat #PI28908).

### Ubiquitination assay and immunoblotting

The ubiquitination assay was performed as described previously (Menon et al., 2015; Boyer et al., 2020). Cells were treated with MG-132 for 1 hour before lysis in buffer containing 20 mM Tris-Cl, 250 mM NaCl, 3 mM EDTA, 3 mM EGTA, 0.5% NP-40, 1% SDS, 2 mM DTT, 5 mM N-ethylmaleimide, protease, and phosphatase inhibitors. Lysates were vortexed, boiled for 20 minutes, cleared at 20,000 x g for ten minutes and diluted in lysis buffer lacking SDS to dilute the final SDS concentration to 0.1%. Myc monoclonal antibody was added and incubated overnight at 4°C with rocking. Protein A beads were added in the morning and incubated for 2 hours. After washing twice with lysis buffer, beads were resuspended in 1x Laemmli sample buffer. Samples were loaded on a 7.5% SDS-PAGE gel, transferred on nitrocellulose membrane for 90 min (75 V), blocked in 5% milk in TBST for 1 hr (RT), and incubated with primary antibody overnight in 1% milk in TBST. Membranes were washed 3x (5 minutes) in TBST, incubated with secondary antibodies for 1 hr, washed 3x (10 minutes) in TBST, rinsed in PBS and imaged on an Odyssey (LI-COR Biosciences).

### Mass spectrometry

HEK293 cells were transfected with Myc-VASP using Lipofectamine-2000. Cells were lysed and spun at 20,000 x g for ten minutes. Lysate was incubated with Myc-Trap beads (Chromotek) for 2 hours. Beads were spun down at 200 x g, washed once with lysis buffer, and washed three times with wash buffer (lysis buffer containing 500 mM NaCl and 1% Triton). After a final wash with 1% SDS in PBS, beads were resuspended in 1x sample buffer (Bio-rad) with 50 mM DTT. Lysis buffer was composed of 50 mM Tris (pH 7.4), 150 mM NaCl, 1 mM EDTA, 0.5% Triton, 150 μg/mL PMSF, 2 μg/mL leupeptin, 5 μg/mL aprotinin, 15 mM sodium pyrophosphate, 50 mM NaF, 40 mM Bgly,1 mM sodium vanadate, 10 mM NEM, 50 μM PR-619.

Immunoprecipitated samples were subjected to SDS-PAGE and stained with coomassie. The bands corresponding to VASP were excised and the proteins were reduced with 10mM DTT for 30 min at RT, alkylated with 100mM iodoacetamide for 45 min in the dark at RT, and in-gel digested with either ArgC (Promega) overnight at 37°C or trypsin (Promega) for 1 hr at 37°C. Peptides were extracted, desalted with C18 spin columns (Pierce) and dried via vacuum centrifugation. Peptide samples were stored at-80°C until further analysis. The peptide samples (n=2) were analyzed in technical duplicate by LC/MS/MS using an Easy nLC 1200 coupled to a QExactive HF mass spectrometer (Thermo Scientific). Samples were injected onto an Easy Spray PepMap C18 column (75 μm id × 25 cm, 2 μm particle size) (Thermo Scientific) and separated over a 45 min method. The gradient for separation consisted of 5–38% mobile phase B at a 250 nl/min flow rate, where mobile phase A was 0.1% formic acid in water and mobile phase B consisted of 0.1% formic acid in 80% ACN. The QExactive HF was operated in data-dependent mode where the 15 most intense precursors were selected for subsequent fragmentation. Resolution for the precursor scan (m/z 350–1600) was set to 60,000 with a target value of 3 × 106 ions. MS/MS scans resolution was set to 15,000 with a target value of 1 × 105 ions, 100 ms injection time. The normalized collision energy was set to 27% for HCD. Dynamic exclusion was set to 30 s, peptide match was set to preferred, and precursors with unknown charge or a charge state of 1 and 8 were excluded.

Raw data files were processed using Proteome Discoverer version 2.5 (Thermo Scientific). Peak lists were searched against the reviewed Uniprot human database, appended with a common contaminants database, using Sequest. The following parameters were used to identify tryptic or ArgC peptides: 50 ppm precursor ion mass tolerance; 0.02 Da product ion mass tolerance; up to 3 missed trypsin cleavage sites; (C) carbamidomethylation was set as a fixed modification; (M) oxidation, (S, T, Y) phosphorylation, and (K) diGly were set as variable modifications. The ptmRS node was used to localize the sites of phosphorylation and diGly sites. Peptide false discovery rates (FDR) were calculated by the Percolator node using a decoy database search and peptides were filtered using a 1% FDR cutoff. MS/MS spectrum was annotated using IPSA (van der Wal et al., 2018). MS Data are available via ProteomeXchange (Perez-Riverol et al., 2022, 2016; Deutsch et al., 2023) with identifier PXD041196

### Protein purification

GST-Ubiquitin (1-74, G75C) was transformed into RIL E. coli cells and grown in LB media with 200 μg/mL of ampicillin and 25 μg/mL of chloramphenicol. Cultures were grown at 37°C until the O.D. reached 0.8-1. IPTG (0.6 mM) was added and cultures were grown overnight at 18°C. Bacteria were pelleted, resuspended in lysis buffer (50mM Tris (pH 8), 200mM NaCl, 1mM DTT, and 2.5mM PMSF), and frozen until use. Pellets were thawed, lysed by Emulsiflex, and centrifuged for 40 minutes (36,600 x g, SS-34 rotor). Protein was bound to Pierce gluthathione agarose (ThermoFisher, 16101) and washed twice with four CV of wash buffer (10mM Tris HCl, pH 8, 250mM NaCl; 1mM DTT). Hrv-3c protease was added at a 1:200 ratio in wash buffer to perform on-bead cleavage of the protein.

His-VASP was transformed into RIL *E. coli* cells and grown in LB media with 200 μg/mL of ampicillin and 25 μg/mL of chloramphenicol. Cultures were grown at 37°C until the O.D. reached 0.8-1. IPTG (0.6 mM) was added and cultures were grown overnight at 18oC. Bac-teria were pelleted, resuspended in lysis buffer (50 mM HEPES pH 7.4, 300 mM NaCl, 10 mM imidazole, 10 mM β-mercaptoethanol), and frozen until use. Pellets were thawed, lysed by sonication, and cleared with a high speed spin. His-VASP was bound to HisPur™ Ni-NTA Resin (ThermoFisher #88221), washed twice with 5 CV of wash buffer (50 mM HEPES pH 7.4, 300 mM NaCl, 30 mM imidazole, 10 mM β-mercaptoethanol), and eluted with 4 CV of 50 mM HEPES pH 7.4, 300 mM NaCl, 250 mM imidazole, 10 mM β-mercaptoethanol. Eluted His-VASP was dialyzed overnight in 20 mM HEPES (pH 7.4), 200 mM NaCl, and 1mM DTT before separation over a Superose 6 column (GE) on an AKTA FPLC (GE).

Actin was purified from chicken skeletal-muscle ace-tone powder (Spudich et al., 1971) and labeled on lysines with Alexa488-succinimidylested (Isambert et al., 1995). Human fascin and mouse capping protein (dual-expression construct of MmCP α1-His(6x)-tag and SNAP-β2; SNAP-CP) were expressed in bacteria and purified as previously described (Vignjevic et al., 2006) (Burke et al., 2017).

### Crosslinking

Crosslinking protocols were adapted from previously published work (Brown et al., 2015). His-VASP constructs and Ubiquitin G75C were reduced with 10 mM DTT and buffer exchanged into reaction buffer (50 mM HEPES (pH 7.0), 200 mM NaCl). Ubiquitin was modified with 10-molar excess BMOE (Fisher) for 30 minutes on ice and desalted into fresh reaction buffer. A 10-fold excess of Ubiquitin-BMOE was added to His-VASP K240C and His-VASP K286C, with a 20-fold excess added to His-VASP K240C, K286C. After a 1-hour incubation (on ice), the reaction was quenched with 10 mM BME. To remove excess (non-crosslinked) ubiquitin, His-VASP was re-purified by nickel affinity chromatography. Protein was buffer exchanged to remove excess imidazole and incubated with His-TEV overnight at 4°C.

### Sortase labeling

A custom peptide (TAMRA-PEG6-LPETGG) was purchased from BioMatik to facilitate fluorescent labeling of VASP. After confirming cleavage of the His-tag from VASP using TEV protease, the sortase reaction was set up with a 20-fold excess of sortase peptide and 2.5 μM His-sortase in a reaction buffer (50 mM HEPES, pH 7.4, 200 mM NaCl, 10 mM CaCl2). After 1 hr incubation (on ice), the reaction was quenched with 20 mM EGTA and buffer-exchanged to remove excess peptide. His-TEV and His-sortase were removed following incubation with nickel resin. TAMRA-VASP ubiquitination mimics were further purified over a Superose 6 column. Purified protein was stored in 10 mM HEPES, 200 mM NaCl, 1 mM DTT, and 10% glycerol, aliquoted, and flash frozen.

### Mass photometry

Measurements were completed on a Refeyn One Mass Photometer (Refeyn Ltd, Oxford, UK) and analyzed using DiscoverMP software (Refeyn Ltd, Oxford, UK) as previously described (S. Cannon et al., 2023). Briefly, imaging wells were assembled from imaging coverslips (24 × 50 mm2, Thorlabs) and adhesive, four-well gaskets (Thorlabs). Measurements were acquired for 1 min at a frame rate of 100 fps at room temperature using AcquireMP software (Refeyn Ltd, Oxford, UK) using 15 nM VASP.

### Actin co-sedimentation assays

Co-sedimentation assays were adapted from previously described protocols (Zimmermann et al., 2016). Lyophilized rabbit skeletal muscle actin (99% pure, Cytoskeleton) was resuspended in water and diluted to 10 mg/mL in fresh Ca-G buffer (2mM Tris, pH 8, 0.2mM ATP, 0.1mM CaCl2, 0.5mM DTT, 0.01% NaN3). After a one hr incubation on ice to depolymerize actin oligomers, actin was centrifuged (20,000 x g, 4°C) for 20 minutes, and the concentration was confirmed. Actin polymerization was induced by the addition of 10 mM HEPES (pH 7.4), 10 mM EGTA, 10 mM MgCl2, 50 mM NaCl, and 1 mM DTT and filaments were incubated for 1 hr at RT.

For low speed sedimentation assays, varying amounts of VASP (100-600 nM) and actin filaments (2 μM) then incubated for 1 hr at RT before centrifugation at 10,000 x g (RT) in a tabletop centrifuge. The top 50% of the supernatant was removed, mixed with 4xSB, and boiled. The remaining 50% of the supernatant was discarded. The pellet was washed carefully in 10 mM HEPES (pH 7.4), 10 mM EGTA, 10 mM MgCl2, 50 mM NaCl, and 1 mM DTT. The pellet was then directly resuspended in 1xSB, transferred to an Eppendorf tube, and boiled. For high speed sedimentation assays, varying amounts of VASP (200-600 nM) and actin filaments (1 μM) then incubated for 1 hr at RT before centrifugation at 100,000 x g (23°C) in a TLA100 rotor (Beckman-Coulter). The top 50% of the supernatant was removed, mixed with 4xSB, and boiled. The remaining 50% of the supernatant was discarded. The pellet was washed carefully in 10 mM HEPES (pH 7.4), 10 mM EGTA, 10 mM MgCl2, 50 mM NaCl, and 1 mM DTT. The pellet was then directly resuspended in 1xSB, transferred to an Eppendorf tube, and boiled. Supernatant and pellet fractions were separated on SDS-PAGE gels and stained with Collodial Coomassie (Invitrogen) for 3 hrs. After destaining in water, gels were scanned on a Licor Odyssey and analyzed by densitometry in ImageStudio.

### Pyrene actin elongation assays

Pyrene elongation assays were adapted from previously described protocols (Zimmermann et al., 2016). Before starting, both rabbit skeletal muscle actin (Cytoskeleton) and pyrene-labeled rabbit skeletal muscle actin (Cytoskeleton) were spun at 100,000 x g for 1 hr at 4 oC a TLA100 rotor (Beckman-Coulter). After conversion of Ca2+-ATP-actin to Mg2+-ATP-actin, polymerization of actin filaments (5 μM) was initiated by adding in 10 mM HEPES (pH 7.4), 10 mM EGTA, 10 mM MgCl2, 50 mM NaCl, and 1 mM DTT and incubated for 1 hour at RT. Additional proteins (VASP and/or capping protein) were added to a final concentration of 0.5 μM filaments before the addition of 0.5 μM actin monomers (20% pyrene-labeled). Pyrene fluorescence was measured with a Spark Cyto (Tecan) or a CLARIOstar (BMG Labtech) plate reader.

### Microscope Descriptions

TIRFM images were acquired using an Olympus IX-71 microscope through TIRF illumination, recorded with a iXon EMCCD camera (Andor Technology), and a cellTIRF 4Line system (Olympus).

All widefield imaging was performed using an inverted microscope (IX83-ZDC2; Evident/Olympus) equipped with a cMOS camera (Orca-Fusion, Hamamatsu) and Xenon light source Images were acquired using a 100× 1.50 NA TIRF objective (Olympus) and with Cellsens software (Evident/Olympus).

Confocal microscopy was performed on an inverted laser scanning confocal microscopy (LSM980, Zeiss) with a motorized X,Y stage and Z focus with high speed Piezo insert. The microscope is equipped with 4 diode lasers (405, 488, 561, 633). Images were acquired with 1x Nyquist sampling using a 63x 1.4 NA Plan-Apochromat oil objective (Zeiss) and 4 channel GaAsP detectors on Zen Blue 3.6 software. This microscope is equipped with Airyscan 2, however, this module was not used in these experiments.

### TIRFM Assays

Microscope slides and coverslips (#1.5; Fisher Scientific, Waltham, MA) were sonicated for 2 h with Helmanex III detergent (Hellma Analytics, Müllheim, Germany). After rinsing in deionized water and drying, glass was incubated with 1 mg/mL mPeg-Silane (5000 MW) in 95% ethanol, pH 2.0 overnight while shaking. Imaging flow chambers were created using double-sided tape to form parallel wells (Zimmermann et al., 2016).

All TIRF assays utilized purified chicken muscle actin. Spontaneous actin assembly was initiated by mixing 1.5 μM Mg-Actin monomers (10% AF488 labeled) with 1x TIRF buffer (10 mM imidazole [pH 7.0], 50 mM KCl, 1 mM MgCl2, 1 mM ethylene glycol tetraacetic acid [EGTA], 50 mM dithiothreitol [DTT], 0.2 mM ATP, 50 μM CaCl2, 15 mM glucose, 20 μg/ml catalase, 100 μg/ml glucose oxidase, and 0.5% [400 cP] methylcellulose), as well as any additional proteins such as VASP and Fascin. Samples were then transferred to flow chambers and imaged at RT by TIRFM.

Actin elongation rates were completed by manually tracking the elongation of single actin filaments or parallel actin bundles in FIJI and the average elongation rate for each filament was then calculated.

Quantification of 1 nM VASP at the barbed ends of actin filaments (in the presence of 673 nM fascin) was adapted from previously published work (Harker et al., 2019). VASP residence was specifically quantified on the trailing filaments of 2-filament bundles. Linescans were used to trace actin filaments and kymographs were created in ImageJ to visualize the growth of the trailing barbed end. Dynamic VASP puncta binding was quantified; stationary puncta were excluded, as they were assumed to be adhered to the coverslip. Likewise, puncta on actin bundles where the trailing versus leading filament was not evident were not included. Cumulative frequency of dwell times were calculated and (1-Cumulative Frequency) was fit with a one-phase decay curve as described previously (Hansen and Mullins, 2010).

### Neon electroporation of MV^D7^ cells

25mm #1.5 german glass coverslips (Electron Microscopy Services Cat #72290-12) ¬were cleaned with a Harrick Plasma Cleaner (PDC-32G). Coverslips were coated with 10 μg/mL Fibronectin (Corning #356008) and incubated for 1 hr at 32°C.

MVD7 were electroporated using the Neon™ Transfection System (ThermoFisher Scientific MPK5000). Briefly, cells were grown to 70-90% confluency, removed from the dish with Accutase (Sigma-Aldrich Cat #A6964), washed with PBS, and resuspended in Neon™ Resuspension Buffer R at a final density of 0.5 × 10^8^ cells/mL. In the final 10 μL Neon reaction, 1.1 μM of protein and 440,000 cells were used. Neon settings were pulse voltage (1250), pulse width (30 ms), pulse number (1).

220,000 cells were plated per coverslip and incubated for 30 minutes at 32°C. Cells were fixed by adding equivolume warm 8% PFA in 2x PHEM buffer (120 mM PIPES, 50 mM

HEPES, 20 mM EGTA, 4 mM MgSO4, and 0.24 M sucrose) directly to the media for 20 min at room temperature. Coverslips were washed 3x with PBS. After fixation, cells were permeabilized with 0.1% TritonX-100 for 10 minutes and blocked with 10% donkey serum (Fisher Scientific Cat #OB003001) for 30 min. To visualize actin filaments, cells were incubated with AlexaFluor-488/647-Phalloidin at 1:400 or 1:200 (respectively) for 1 hour at room temperature. Coverslips were washed 3x with PBS, mounted in a homemade Tris/glycerol/n-propyl-gallate–based mounting media and sealed with nail polish.

### MV^D7^ cell analysis

All images were processed and analyzed in ImageJ (Schindelin et al., 2012). For classification of widefield cell spreading images, images were blinded. Cell masks were created using the phalloidin channel to quantify cell area, perimeter, and fluorescence intensity. Cell classification was based on the presence of a smooth and even lamellipodia (smooth), a filopodial density greater than 0.1 filopodia/um perimeter (filopodial), and the presence of actin ruffles (ruffled) as previously described (Applewhite et al., 2007).

Maximum projections were created from confocal images to analyze lamellipodia and filopodia protein enrichment. Images were manually background subtracted using an average of blank regions on the coverslip. GFP-VASP images were used to draw linescans on each structure. To account for variability in lamellipodia thickness, four linescans were drawn per cell in areas with clear lamellipodia (per GFP-VASP localization). Filopodia linescans were drawn from the filopodia tip to the base.

Raw filopodial tip intensity values were calculated by averaging the first four pixels (0.255 μm) of the filopodia. Heatmaps displaying protein localization were created based on FiloMap code (Jacquemet, 2023).

### Software

All data analysis was completed in Prism 9 (GraphPad). Figures were created in Adobe Illustrator.

## Supplementary Information

Fig S1 displays MS/MS plots for ubiquitinated VASP peptides. Fig S2 shows filopodia number (in filopodial cells), cell area, and phalloidin intensity is unchanged with TAMRA-VASP or TAMRA-VASP mmUb-K240/K286 electroporation into MVD7 cells. Video 1 and 2 shows actin polymerization in the presence of VASP or ubiquitinated VASP corresponding to Fig 3D,E and Fig 3F, respectively. Table S1 details the curve fitting parameters from seeded actin pyrene assays and Table S2 details curve fitting parameters from seeded actin pyrene assays in the presence of capping protein

## Acknowledgements

This work was supported by National Institutes of Health Grants R35GM135160 (S.L.G.), 1F31NS113381 (L.E.M), R35GM128855 (N.G.B.), and R01GM079265 (D.R.K).

Mass spectrometry data was generated using the UNC Proteomics Core Facility, which is supported in part by NCI Center Core Support Grant (2P30CA016086-45) to the UNC Lineberger Comprehensive Cancer Center. Mass photometry was performed at the UNC Center for Structural Biology, which is supported by the National Cancer Institute of the National Institutes of Health under award number P30CA016086. Confocal microscopy was performed at the UNC Neuroscience Microscopy Core, which receives funding from the NIH-NINDS Neuroscience Center Support Grant P30 NS045892 and the NIH-NICHD Intellectual and Developmental Disabilities Research Center Support Grant P50 HD103573. In particular, this work utilized the LSM980 microscope at the Neuroscience Microscopy Core, which was funded with support from NIH grant S10 OD032388. The authors declare no competing financial interests.

