## Supplemental Materials for "Multi-monoubiquitination controls VASP-mediated actin dynamics"

### Supplemental Material

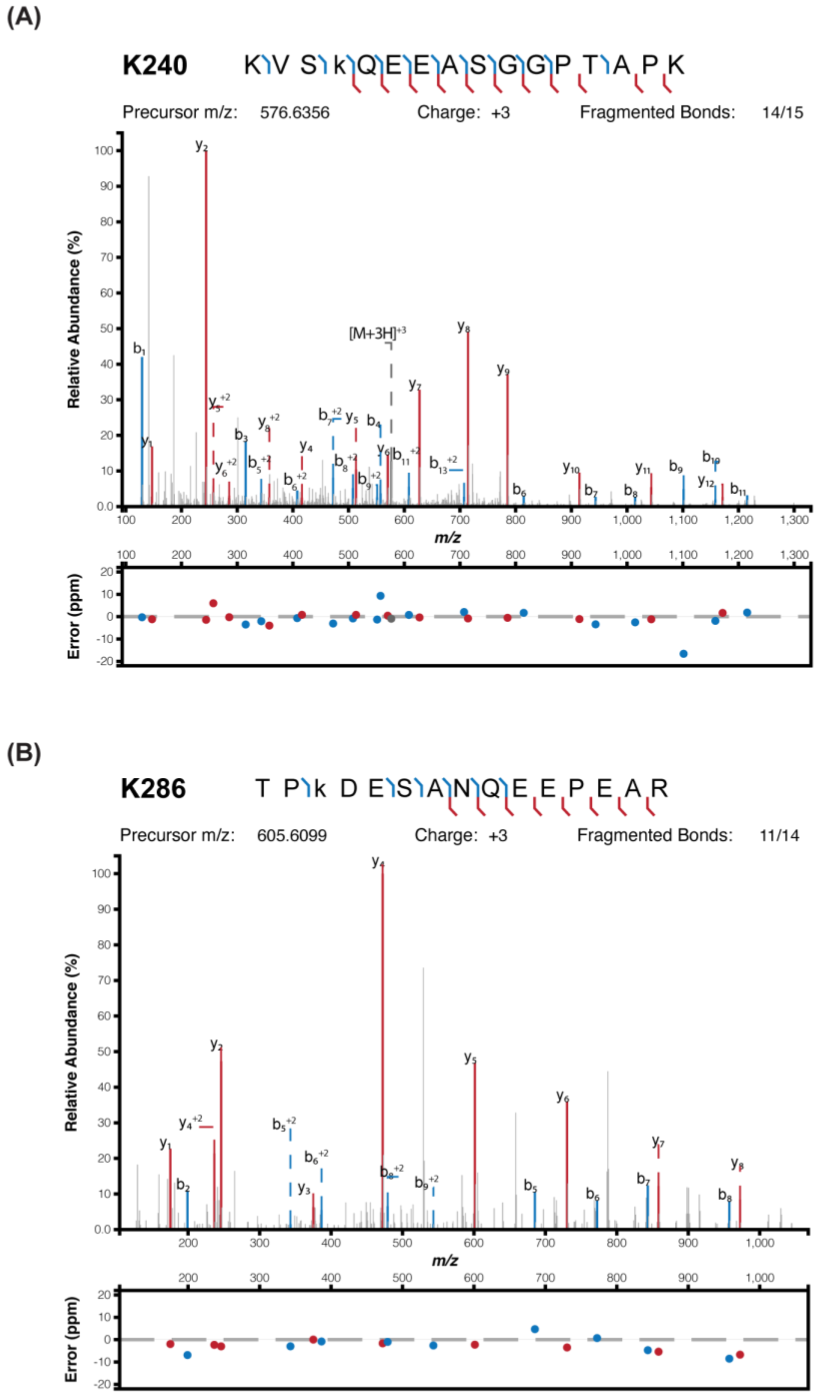

**Fig S1. MS/MS Spectra of ubiquitinated peptides.**

(A) MS/MS spectrum of the triply charged ion (m/z 576.9678) corresponding to VASP tryptic peptide KVSKQEEASGGPTAPK. K4 is diGly modified (k), corresponding to K240 in VASP.

(B) MS/MS spectrum of the triply charged ion (m/z 605.6099) corresponding to VASP tryptic peptide TPKDESANQEEPEAR. K3 is diGly modified (k), corresponding to K286 in VASP

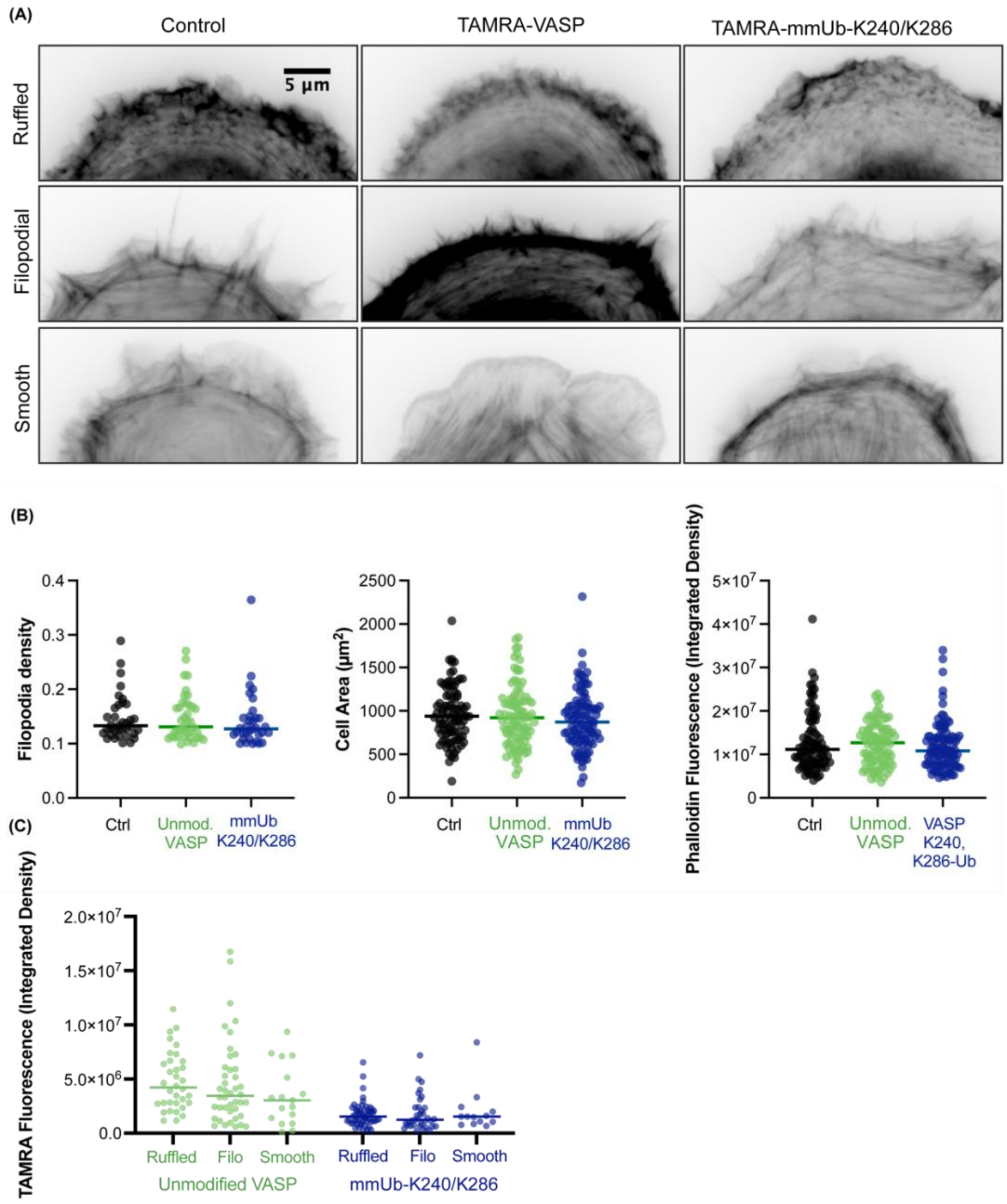

**Figure S2. Electroporation of mmUb-K240/K286 does not change filopodial density, cell area or phalloidin-stained actin levels.**

(A) Representative widefield phalloidin images of each cell spreading phenotype.

(B) Quantification of filopodia density, cell area, and phalloidin fluorescence intensity. 92-104 cells were quantified for each condition across three experiments.

(C) TAMRA fluorescence intensity separated by cell spreading phenotype. N = 13-54 cells per classification across three experiments.

| | Unmodified | mUb-K240 | mUb-K286 | mUb-K240/K286 |
|---|---|---|---|---|
| **Bmax** | 45.76 | 31.66 | 41.86 | 17.71 |
| **95% CI** | 37.62 to 57.65 | 22.25 to 48.8 | 21.92 to +∞ | 7.533 to +∞ |
| **$K_D$ (nM)** | 25.45 | 17.99 | 43.75 | 54.89 |
| **95% CI** | 14.24 to 46.30 | 5.347 to 58.12 | 5.932 to +∞ | 5.122 to +∞ |

**Table 1. Curve-fitting parameters for actin pyrene elongation assays.**

| | Unmodified | mUb-K240 | mUb-K286 | mUb-K240/K286 |
|---|---|---|---|---|
| **Bmax** | 33.56 | 28.55 | 33.93 | 27.04 |
| **95% CI** | 27.59 to 41.15 | 23.61 to 34.58 | 23.61 to 34.58 | 28.17 to 41.17 |
| **$K_D$ (nM)** | 37.83 | 24.88 | 36.19 | 42.88 |
| **95% CI** | 18.18 to 71.54 | 10.60 to 49.18 | 17.74 to 67.02 | 21.43 to 78.37 |

**Table 2. Curve-fitting parameters for actin pyrene elongation assays in the presence of capping protein.**

**Video 1 Legend**
Polymerization of 1.5 µM actin (10% AlexaFluor-488 labeled, shown in green) visualized with TIRF microscopy for ten minutes (2.5 s acquisition time) in the presence of 25 nM TAMRA-VASP (shown in magenta). Frame rate 30 fps.

**Video 2 Legend**
Polymerization of 1.5 µM actin (10% AlexaFluor-488 labeled) visualized with TIRF microscopy for ten minutes (2.5 s acquisition time) in the presence of 1.25-10 nM TAMRA-VASP (not shown). Frame rate 30 fps.
